# Farmers’ perceptions on the causes of cassava root bitterness in konzo-affected areas of Mtwara Region, Tanzania

**DOI:** 10.1101/397018

**Authors:** Matema L.E. Imakumbili, Ernest Semu, Johnson M.R. Semoka, Adebayo Abass, Geoffrey Mkamilo

**Affiliations:** Department of Soils and Geological Sciences, Sokoine University of Agriculture, Morogoro, Tanzania; The International Institute of Tropical Agriculture (IITA), Eastern Africa Hub, Dar es Salaam, Tanzania; Naliendele Agricultural Research Institute, Mtwara, Tanzania

## Abstract

The agronomic factors influencing increased cyanogenic glucoside levels, particularly in bitter cassava varieties during periods without water stress, in areas where konzo (a cassava cyanide related paralytic disorder also called spastic paraparesis) persists, are hardly known. However, through their assessment of bitter taste, farmers may have noticed factors unrelated to water stress and variety type that additionally influence cassava root cyanogenic glucoside content in these environments. Bitterness in cassava is usually associated with high cyanogenic glucoside levels. Using some konzo-affected areas in Mtwara region of Tanzania as a case study, a survey was thus carried out to identify the factors, hitherto overlooked, that may additionally influence cyanogenic glucoside levels in cassava. A total of 120 farmers were interviewed. A number of factors unrelated to water stress and variety type that could be additionally influencing cyanogenic glucoside production in cassava plants were mentioned. The mentioned factors included nutrient poor soils, plant age at harvest, weeds, piecemeal harvesting, and branch pruning; the factors, respectively, constituted 14.2%, 7.5%, 0.8%, 0.8%, and 0.8% of the total responses given. The revealed factors constitute permanent environmental characteristics and commonly used crop management practices by farmers living in konzo-prone Mtwara region of Tanzania that could be additionally resulting in high cyanogenic glucoside levels in cassava, regardless of water stress.

## Introduction

Cassava (*Manihot esculenta* Crantz) is the seventh most important food crop in the world [1]. Despite its importance, cassava unfortunately contains potentially harmful cyanogenic glucosides, which release toxic hydrogen cyanide upon hydrolysis [2]. Without access to foods containing sulphur amino acids, continuous ingestion of cyanogenic glucosides from improperly processed cassava products, often results in cases of cassava cyanide intoxication in rural poor cassava dependent communities [3]. Cases of cassava cyanide intoxication have been reported in a number of countries in Sub-Saharan African, such as the Democratic Republic of Congo (DRC), Mozambique, Tanzania, Cameroon, Central African Republic and Angola [4–8]. The reports consisted of cases of acute cyanide intoxication but more commonly of the cassava cyanide health disorder called konzo (spastic paraparesis), which is known to result in an irreversible paralysis of legs [9–12].

One reason given for high cyanide intake during konzo epidemics are increased cyanogenic glucoside levels in fresh cassava roots and in cassava products produced from them [6,13]. Researchers mostly attribute the increase in cyanogenic glucoside levels in cassava plants to water stress from prolonged droughts that coincide with most epidemics of konzo [6,14] and also to the bitter varieties which are preferred by many farmers [6,15]. The naturally high cyanogenic glucoside contents of bitter cassava varieties are said to be further increased by drought [13]. Water stress during dry seasons (seasonal droughts) is similarly known to result in increased cyanogenic glucoside levels of cassava plants [5,16,17]. Dry season water stress is able to increase cassava cyanogen levels by 9 - 10 times the levels obtained during the rainy season [18].

Konzo is persistent in some rural areas of Mozambique, DRC and Tanzania [8]. Small epidemics and sporadic cases of konzo have been concurrently observed in some communities, creating near persistent konzo exposures [7,19]. In areas where konzo persists, high cyanogenic glucoside levels in cassava plants may also occur outside periods of water stress [19]. This suggests that factors other than droughts and dry seasons could also be contributing to increased cyanogenic glucoside concentrations in cassava plants, which may be contributing to the occurrence of persistent cassava cyanide intoxication, or konzo, in these areas. Other agronomic factors that are a permanent characteristic of these farming systems, could thus be additionally influencing cassava cyanogenic glucoside levels.

Farmers in Africa generally use the bitter taste of cassava roots to perceive the potential toxicity of cassava [15,20,21]. Research has also shown that a greater portion of cassava varieties perceived as bitter by farmers do indeed contain higher levels of cyanogenic glucoside than the safer to consume sweet cassava varieties [20]. This may however not always be the case as cassava contains other bitter compounds [22], making validation necessary. Cassava root taste can also be used to assess changes in the degree of bitterness of roots of a particular cassava variety [23,24]. During a season in which a konzo epidemic was experienced, families had been reported to complain that cassava roots were more bitter than normal [25]. Through taste perceptions, most people had become conscious of an increase in cassava root bitterness that led to its associated toxicity. This shows how the bitter taste of cassava roots can be used to determine increased bitterness.

When sporadic cases of konzo occur un-related to water stress, it is more difficult to explain the agronomic factors leading to increased cyanogenic glucoside levels, particularly in the bitter cassava varieties. However, being able to observe the crop throughout the year, farmers may have knowledge of other agronomic factors influencing cassava root bitterness and consequently root cyanogenic glucoside levels. Hence, using some konzo-affected communities in Mtwara region, in Tanzania, this research was carried out to investigate agronomic factors, other than variety and water stress that could be additionally influencing cassava root bitterness, according to the perception of farmers.

## Materials and methods

### Description of the study area

Three districts have been reportedly affected by cassava cyanide poisoning in Mtwara region, namely Masasi, Mtwara Rural and Newala districts [6,9,26]. This study, however, focused on villages of Mtwara Rural and Newala districts. The two districts covered in the survey are two of the five districts found in Mtwara region (S 10^°^16’25”, E 40^°^10’58”). The soils in the Mtwara region and in the two districts (Mtwara and Newala) have low natural soil fertility [27,28]. They are predominantly sandy and have been classified as Ferralic Cambisols [27,29]. The study areas mainly lie in Tanzania’s Coastal Lowlands agroecological zone (C 2) [27,29]. The rainfall in the study area is mono-modal and ranges from 800 to1000 mm/year and the maximum and minimum temperatures vary from 29 to 31 °C and between 19 to 23 °C, respectively [27].

### Sampling method

In October 2014, a total of 120 household heads (male or female) with full knowledge of the cropping history of their farm fields were selected to be interviewed from eight randomly selected konzo-affected villages of Newala and Mtwara Rural districts. Sixty-one of the respondents were from Newala district and 59 were from Mtwara Rural district. The villages selected were among the 18 villages visited during a konzo rehabilitation and prevention program that was carried out, from 2008 to 2009, through collaboration between the Tanzania Food and Nutrition Centre (TFNC) and the Tanzania Red Cross Society (TRCS), with technical support from Australian National University and with funding from AusAID [6]. Using the 2012 census list, 15 households from each village were randomly picked for interviews. Each household was first assigned a unique number and the first 15 households were picked after randomising the numbered village census list using the ‘RAND’ function in Microsoft Excel.

### Field methods and tools

A questionnaire containing both closed and open ended questions was used to collect information on what farmers perceived to be the causes of increased bitterness of cassava roots. Open ended questions were used to allow the farmer to provide further explanations to closed ended responses. Visits to the fields were also made to observe how the households practice cassava cultivation.

### Data Analysis

The collected data was analysed as frequencies, using GenStat package, Edition 14. Each agronomic factor mentioned had 120 chances of being mentioned, according to the total number of respondents. The sum of responses given for a mentioned factor was determined and frequency percentages were then calculated based on the maximum number of responses that could be given.

## Results and Discussion

Table 1 shows the factors perceived by farmers as contributors of cassava root bitterness. The percentage of farmers that attributed root bitterness to each factor is also shown. The factors can be categorised into genotype (variety), environment and crop management factors. Some factors were mentioned more than others but this does not mean that they are less significant contributors of cassava root bitter, this can only be proved by research.

**Table 1.** Farmers’ perceptions (responses) on factors influencing bitterness in cassava roots

| Factors influencing root bitterness | Mtwara Rural District |  | Newala District |  | Total (both districts) |  |
| --- | --- | --- | --- | --- | --- | --- |
|  | n = 59 | (%) | n = 61 | (%) | n = 120 | (%) |
| 1. Variety | 59 | 100.0 | 61 | 100.0 | 120 | 100.0 |
| 2. Environmental factors |  |  |  |  |  |  |
| a. Soil characteristics | 1 | 1.7 | 16 | 26.2 | 17 | 14.2 |
| b. Seasons (wet or dry) | 5 | 8.5 | 0 | 0.0 | 5 | 4.2 |
| 3. Farmers' agronomic practices |  |  |  |  |  |  |
| a. Plant age at harvest | 0 | 0.0 | 9 | 14.8 | 9 | 7.5 |
| b. Poor weeding | 1 | 1.7 | 1 | 1.6 | 2 | 0.8 |
| c. Piecemeal harvesting | 1 | 1.7 | 0 | 0.0 | 1 | 0.8 |
| d. Branch pruning | 1 | 1.7 | 0 | 0.0 | 1 | 0.8 |

### Variety

Farmers described variety as being the ultimate contributor to cassava root bitterness. They explained that if a variety was bitter then the roots it would produce would be naturally bitter and vice versa for sweet varieties. In agreement with the farmers’, researchers also attribute the cyanogenic character of a variety as being due to an inheritable trait involving a specific gene [30,31]. The production of small amounts of cyanogenic glucosides in sweet varieties is regulated by a recessive gene complex [32].

## Environmental factors

### Soil characteristics

Soil type was perceived as a contributing factor to cassava bitterness by 17 (14.2%) respondents, with most of them being from Newala district (Table 1). It was the second most perceived contributor of cassava root bitterness. Although most farmers were often not specific about which cassava varieties changed taste with soil type, the variety *Kigoma* had been specifically identified as having the tendency of changing from sweet to bitter with soil type. Cassava bitterness was associated with four soil factors or types (Fig 1).

**Fig 1.**
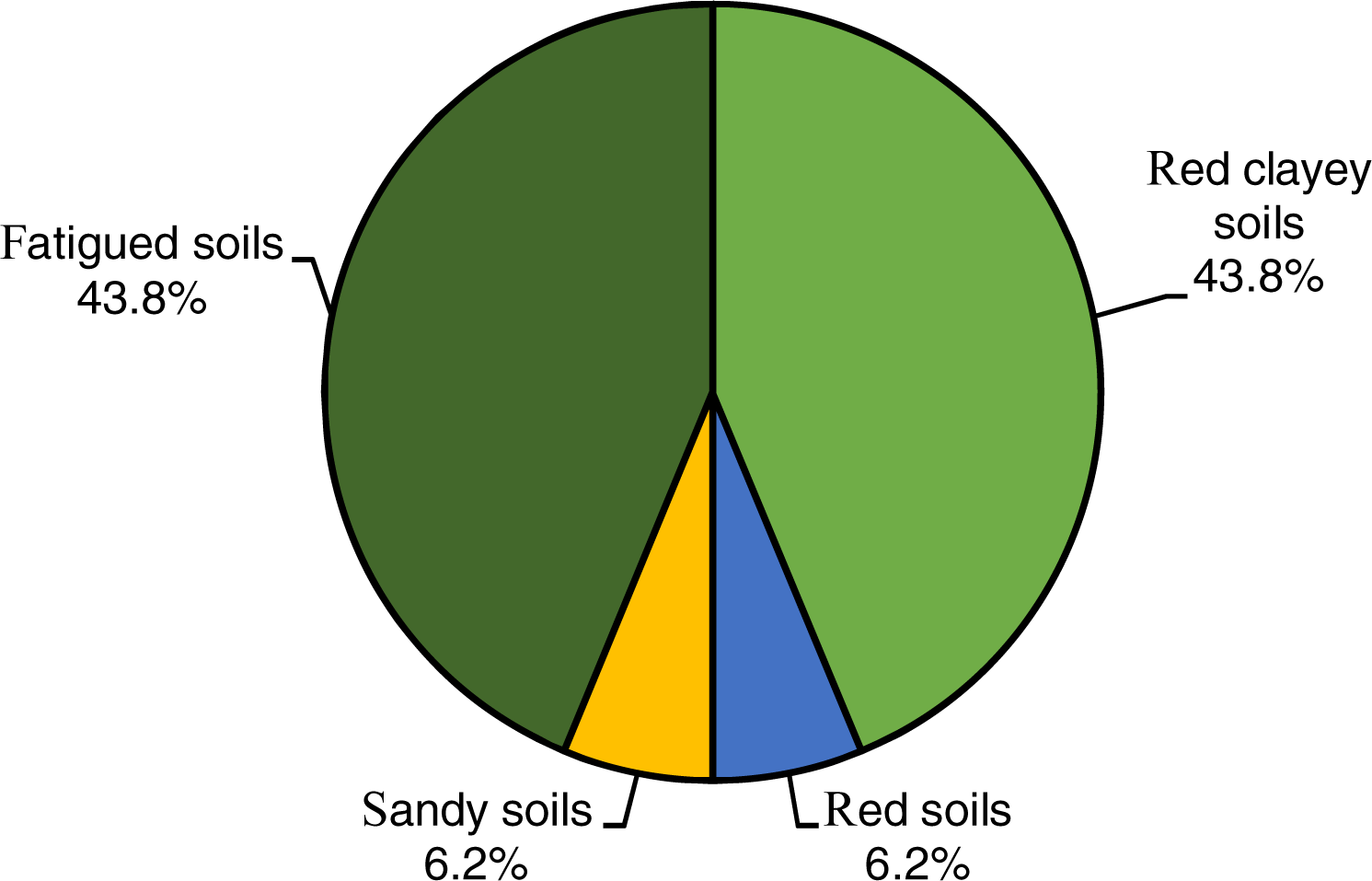
Farmers’ responses on soil factors or types associated with bitterness in cassava roots.

Fatigued soils and red clayey soils were identified by the farmers as the major causes of bitterness in cassava roots. Farmers described fatigued soils as soils that had lost their fertility due to being continuously cultivated. They described red clayey soils as being red coloured soils that could be used for building. Fewer respondents mentioned red soils (6.2%) and sandy soils (6.2%). It is not clear whether the red soils mentioned by some respondents are the same as the red-clayey soils mentioned by others. The two categories of ‘red’ soils were thus recorded separately here. However, when combined, 50.0% of the respondents considered the red soils as problematic at causing cassava root bitterness.

Farmers gave special emphasis to the red-clayey soils located in the Makote area of Newala district. These soils were described as having a thin ‘white’ top soil layer which was then followed by a thick red coloured soil layer from which it got its description. The majority of soils on the Makonde plateau have a thin organic layer overlying a subsoil which increases in redness with depth [33]. Although paler coloured soils are also present, red soils are more common on the Makonde plateau [33]. The red soils are described as being impoverished and as having a poor nutrient holding capacity due to their low cation exchange capacity (< 10 cmol/kg) [33]. The red colour of soils indicates higher concentrations of red coloured manganese dioxide and unhydrated iron oxide minerals [34,35]. High concentrations of iron and manganese oxide minerals indicate a high degree of weathering in soils [36]. The red soils thus contain higher proportions of nutrient poor minerals, making them low in fertility. The farmers believed that red soils change the taste of roots of sweet cassava varieties to bitter. Some farmers mentioned that red soil under anthills is what caused cassava grown on these soils to become bitter.

Some farmers believed that some cropping fields easily became ‘tired’ or ‘fatigued’ after being cultivated for only a few years (Fig 1). They attributed the fatigue or exhaustion of cropping fields to loss of soil fertility. Soil fatigue is defined as the exhaustion of the soil through depletion of essential plant nutrients [37]. Hence, like the red soils, fatigued soils are also nutrient poor. Low soil fertility can cause nutrient stress in cassava [38,39] and probably increase root bitterness and cyanogenic glucoside levels. The short fallow periods (4 years on average) practiced in both districts [40] do not allow for proper soil nutrient and carbon restoration [41], which further enhances nutrient depletion.

A few studies have been carried out to demonstrate the effects of poor soil fertility on cassava root bitterness or on cyanogenic glucoside accumulation in cassava roots. Some studies have shown that an improved supply of the nutrients nitrogen (N), phosphorus (P) and potassium (K) in soils, could help reduce root total cyanide levels compared [24,42]. Reduced cassava root bitterness has also been reported with an improved supply of N, P and K in soil [24]. These findings probably explain the perceived increase in root bitterness on nutrient poor soils. Conversely, higher total root cyanide levels were obtained in cassava grown on more fertile Fluvisols and Andosols as compared to levels found in cassava grown on the highly weathered nutrient poor Nitisols [43]. Increased cassava root bitterness was also observed in cassava grown on nutrient rich soils with higher levels of basic cations (pH as high as 7.8; K as high as 1.9 cmol/kg) and soil organic matter (as high as 8.2%) [44]. Poor soil fertility may thus not always lead to increased root bitterness.

Sandy soils had additionally been pointed out by a few farmers as a cause of cassava root bitterness (Fig 1). It is important to mention that due to their low CEC and low water holding capacity, the problems associated with sandy soils arise from both poor fertility and low moisture retention [45]. The observed increase in cassava root bitterness on sandy soils could thus be due to nutrient stress in addition to water stress.

### Natural water stress conditions

Five interviewed farmers had observed that natural water stress conditions due to seasonal changes (dry and wet seasons) in the region, constituted important factors that influenced bitterness in cassava roots (Table 1). All the five farmers were from Mtwara Rural district. Some of the farmers were able to mention the varieties that had their root taste changed with season; they were the varieties, *Liwoyoka, Kigoma mafia, Musa Saidi, Nachinyanya, Badi, Vincenti, Mnalile Kuchumba* and *Mtukane*. All the varieties mentioned, except for *Musa Saidi*, were sweet varieties. Most respondents mentioned that the bitter taste arose in the dry season. As previously mentioned, water stress caused by seasonal dry periods is able to increase cyanogenic glucoside levels in cassava roots [13,18,46,47]. Mtwara region experiences long inter-seasonal dry spells (normal dry seasons). Like the farmers’ observations konzo (and thus high cyanogenic glucoside levels in cassava) has been observed to occur in a seasonal pattern, with most cases occurring during the dry seasons [5,16]. One study had found a strong link (r^2^ = 0.9) between normal yearly cyclic changes in precipitation and cassava cyanide intoxication that had resulted in konzo [17]. The findings from other studies support the farmers’ observation of increased cassava root bitterness during the dry season period.

### Farmers’ agronomic practices

After soil type, cassava plant age at harvest or the length of time that roots of matured cassava plants are left un-harvested was the next most stated contributing factor to cassava bitterness (Table 1). The farmers had a tendency of leaving cassava unharvested for long, even if it had reached maturity. The two sweet cassava varieties, *Kigoma* and *Kifuru*, were pointed out as being particularly prone to becoming bitter when left unharvested for long. Cyanogenic glucoside production in cassava is known to be also dependent on growth stage (plant age) [48,49]. A reduction in cyanogen levels was observed with increased age (six, eight, 10 and 12 MAP) in flour produced from roots of a bitter cassava variety [50]. Conversely, no differences were observed in the cyanogen content of fresh cassava roots of a sweet cassava variety harvested at three month intervals, starting from 12 to 24 MAP [46]. Similarly, no significant differences were observed in root cyanogen levels of root parenchyma of a bitter cassava variety at different growth stages [47]. Unlike the farmers’ observations, none of the research findings reported an increase in root bitterness in sweet cassava varieties with plant age. The perceived increase in root bitterness with plant age for sweet cassava varieties may however probably be due to a loss of root quality [51].

The farmers additionally believed that roots of late maturing bitter cassava varieties, when harvested ‘early’ (at 12 – 18 months after planting (MAP)), were immature and very ‘toxic’. They explained that cassava flour produced from roots of young bitter cassava varieties was toxic, even when produced using their more efficient traditional forms of processing. In agreement with the farmers young cyanogenic plants are known to contain higher amounts of cyanogenic glucosides [52]. Leaving bitter cassava varieties to grow for a longer time period, was thus a way of reducing cassava cyanide toxicity. The bitter varieties were usually left unharvested until 24 - 36 MAP, while the sweet varieties were usually harvested at 12 - 18 MAP because they matured early. Due to their ability to bulk early, sweet cassava varieties are also harvested early in other parts of Africa, while because they bulk late, bitter varieties can be left unharvested for up to 39 MAP, but this is also because they store longer when left unharvested in the field [53]. Not all bitter varieties were, however, late maturing. For example, the variety *Mohammed Mfuame*, commonly grown in Mtwara Rural district, was a bitter cassava variety that matured within a year and was thus harvested early. Its bitter roots were preferred as a deterrent to pest while its early bulking was equally appreciated as roots could be quickly obtained without having to wait so long.

Weeds were also a concern to farmers as an additional factor contributing to cassava bitterness. Some farmers explained that when cassava was poorly weeded, the roots it produced tended to be bitter. Weeds tend to grow faster and to seriously compete with cassava for light, water and nutrients, subjecting it to plant stress. Biotic stress factors are able to influence cyanogenic glucoside production in cassava plants [54]. In agreement with the farmers’ perception, delayed weeding or no weeding at maturity had been found to result in increased cassava root bitterness [44].

Some farmers carried out piecemeal harvesting, although it was more common for them to harvest all the cassava in a field all at once. Piecemeal harvesting is a traditional cassava harvesting method used by farmers to achieve longer storage of cassava roots in the field. Piecemeal harvesting involves harvesting only the roots needed at a time, while leaving the rest un-harvested and still attached to the cassava plants in the field [55]. Farmers noted that once the first roots had been removed from a cassava plant, the unharvested roots, still attached to the plant eventually became bitter. Hardly any studies have been done on the effects of piecemeal harvesting on cyanogenic glucoside production. It can however be assumed that just like actively growing young cyanogenic plants or plant parts [52], the now rapidly developing smaller roots left behind after piecemeal harvesting [56,57] may also accumulate high cyanogenic glucoside levels. It is however unclear whether increased root bitterness is observed immediately after harvest or after some days, weeks or months. Furthermore, plants that are piecemeal harvested may go on to form new roots before being harvested again [56] and could thus have high cyanogen levels while still young and actively developing.

A common agronomic practice among cassava farmers in Southern Tanzania is branch pruning (debranching) of cassava plants, while leaving them still growing in the field. It is important to note that debranching is different from ratooning, which involves the cutting off of cassava stems to leave behind only a stump of the cassava plant. Branch pruning was mentioned by one farmer to contribute to bitterness in cassava roots (Table 1). One reason why farmer’s branch pruned their cassava plants was to shorten cassava plants to make browsing easier for goats. This prevented goats from damaging the cassava plants, as goats tended to damage tall plants as they struggled to reach for the leaves. Cassava leaves were hence an important source of ruminant feed in these areas in addition to them being an important source of vegetables. Secondly, cassava branches were sometimes cut from the plant and used as a mat for drying large cassava chips (locally called *makopa*) in the field (Fig 2). This helped avoid placing freshly peeled clean cassava chips on the bare ground (Fig 3).

**Fig 2.**
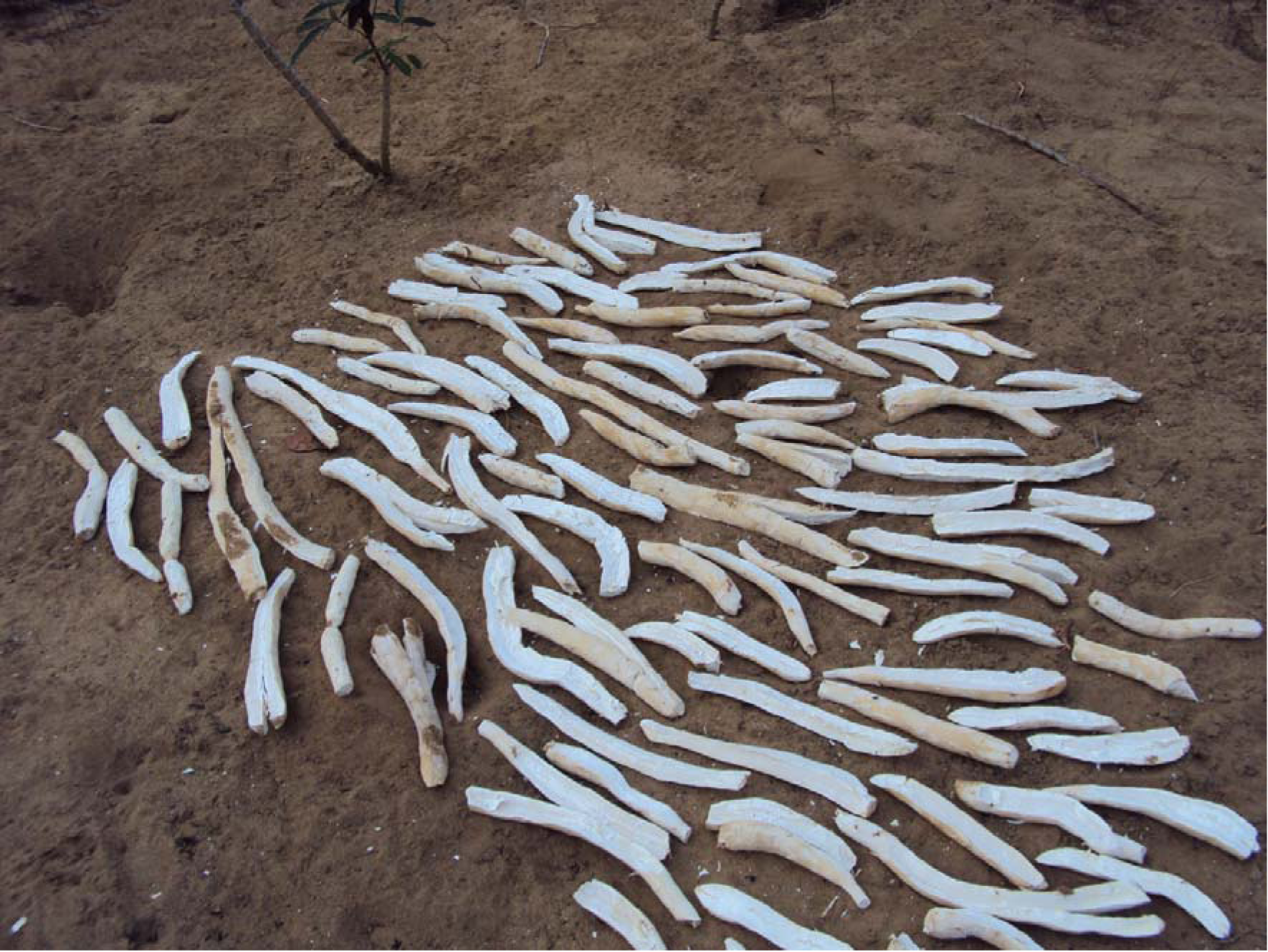
Cassava chips left to dry on cassava stems in the field.

**Fig 3.**
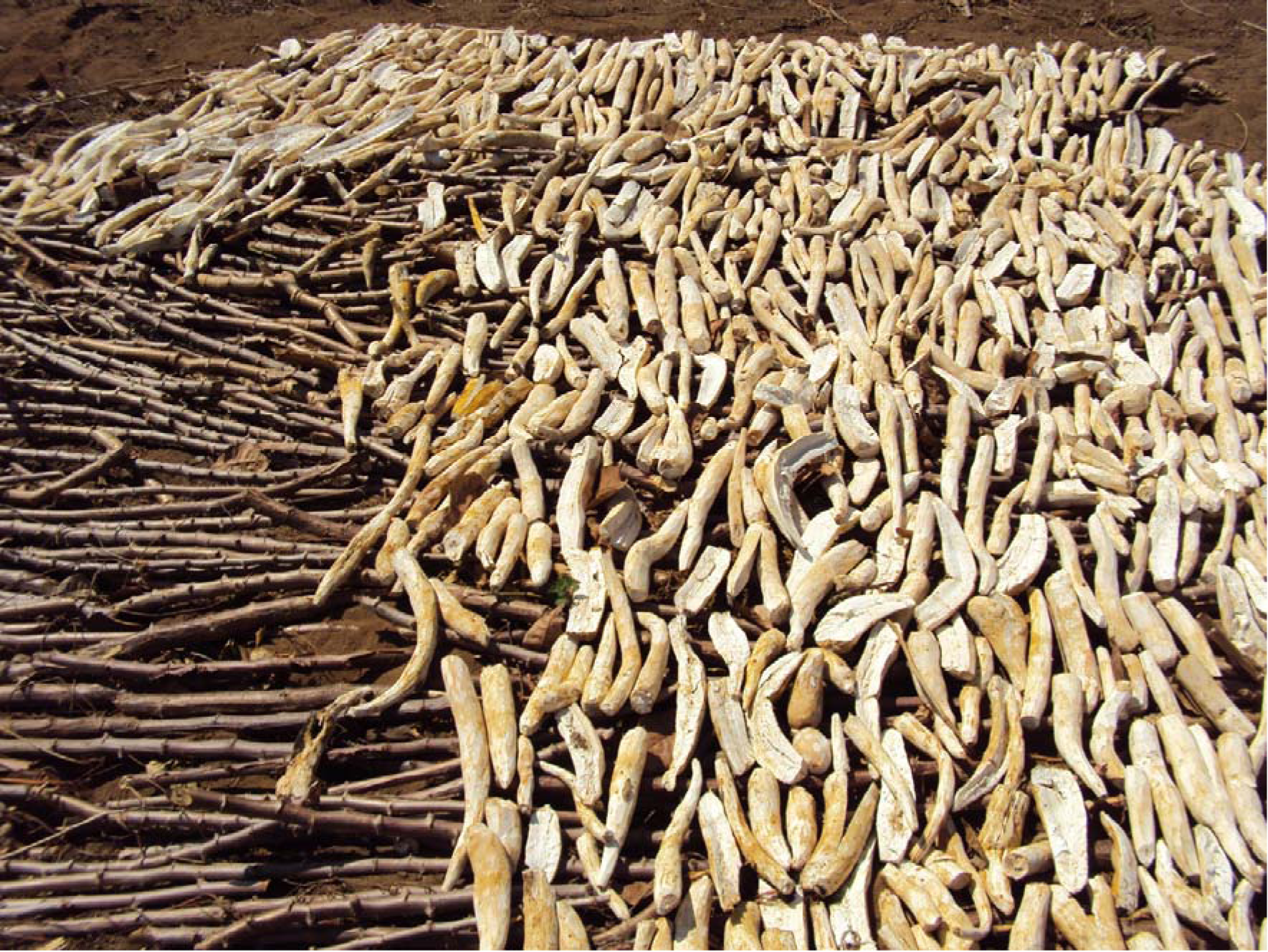
Cassava chips left to dry on bare ground in the field.

There could be other reasons why cassava plants were branch pruned by the farmers. With hardly any research carried out on the effects of branch pruning on cassava root bitterness or on cyanogenic glucoside accumulation, it is difficult to explain the farmers’ observation. However, after being pruned there is a rapid regrowth of cassava shoots and leaves. Since regrowth contains higher levels of cyanogenic glucosides [52], it is possible that more cyanogenic glucosides are transported to roots from the shoots, following pruning. This is because cyanogenic glucosides are synthesised in the leaves of cassava plants and transported to the roots [54,58]. This possibly leads to the root bitterness observed by farmers after cassava plants have been branch pruned.

## Conclusions

The study revealed a number of non-water stress and variety-related factors that according to farmers’ perceptions contribute to bitterness in cassava roots; they include factors such as soil type (especially nutrient low red soils), farmers’ agronomic practices like cassava plant age at harvest, poor weeding, piecemeal harvesting, and branch pruning. These factors may thus be additionally influencing cyanogenic glucoside production in cassava grown in the region. The revealed factors could have contributed to the reported persistent episodes of konzo in Tanzania and could also be contributing to newly occurring cases of konzo in these areas, although not being reported.

The revealed agronomic factors constitute permanent environmental characteristics and practices commonly used by farmers living in konzo-prone areas of Tanzania’s Mtwara region that could be additionally leading to high cyanogenic glucoside levels in cassava roots independent of periods of water stress. Research to validate the effects of the revealed agronomic factors on root cyanogenic glucoside accumulation should be carried out in order to understand their significance. For it to be representative and applicable, the research carried out should however exactly mimic the environment and practices used by farmers in the konzo-prone areas. Lastly, although there are many solutions to tackle cassava cyanide intoxication, agronomic solutions have an important role in making cassava safe to consume right at harvest.

## Acknowledgements

The authors thank the Alliance for a Green Revolution in Africa (AGRA) for making the research possible. They also thank Naliendele Agricultural Research Institute (NARI) for their support during the execution of the research. The authors also thank the farmers that willingly participated in the research.

